# Dendrite injury, but not axon injury, triggers neuroprotection in Drosophila models of neurodegenerative disease

**DOI:** 10.1101/2024.03.29.587363

**Authors:** Sydney E. Prange, Isha N. Bhakta, Daria Sysoeva, Grace E. Jean, Anjali Madisetti, Ly U. Duong, Patrick T. Hwu, Jaela G. Melton, Hieu H. N. Le, Katherine L. Thompson-Peer

## Abstract

Dendrite defects and loss are early cellular alterations observed across neurodegenerative diseases that play a role in early disease pathogenesis. Dendrite degeneration can be modeled by expressing pathogenic polyglutamine disease transgenes in *Drosophila* neurons *in vivo*. Here, we show that we can protect against dendrite loss in neurons modeling neurodegenerative polyglutamine diseases through injury to a single primary dendrite branch. We find that this neuroprotection is specific to injury-induced activation of dendrite regeneration: neither injury to the axon nor injury just to surrounding tissues induces this response. We show that the mechanism of this regenerative response is stabilization of the actin (but not microtubule) cytoskeleton. We also demonstrate that this regenerative response may extend to other neurodegenerative diseases. Together, we provide evidence that activating dendrite regeneration pathways has the potential to slow–or even reverse–dendrite loss in neurodegenerative disease.

## INTRODUCTION

Neurons are the fundamental units of the nervous system and are responsible for receiving and transmitting information throughout the body. To perform this critical function, neurons use dendrites to receive information from the environment and other neurons and axons to send information. Across many neurodegenerative diseases, including Huntington’s disease (HD), amyotrophic lateral sclerosis (ALS), Alzheimer’s disease (AD), Parkinson’s disease (PD), and spinocerebellar ataxias (SCAs), neurons exhibit a number of dendrite defects, including dendrite branch loss, branch thinning, branch shortening, and spine loss during early disease stages and are likely detrimental for neuronal function.^1–7^ For example, in HD mice models, dendritic alterations are observed in pre-symptomatic and symptomatic mice before neuronal death.^4,8,9^ Similarly, in mouse models of ALS, pathological dendrite changes occur before motor deficits in some neuron types and in early or mid-disease stages in other neuron types.^3,10–12^ Because these dendrite defects happen in early disease stages, sometimes pre-symptomatically in animal models, and also appear long before any mass neuronal death associated with late stages of disease, dendrite defects have emerged as a significant contributor to early disease pathogenesis.^6,13^ Dendrite defects are therefore a crucial neuronal alteration preceding cell death that may contribute to the initiation of symptoms in the early stages of neurodegenerative disease. Ameliorating these dendrite defects may provide an opportunity for early intervention and disease-modifying therapies in these conditions.

Dendrites are lost in neurodegenerative disease, but neurons have the capacity to regenerate dendrites. Following severing of all dendrite branches, neurons in the *Drosophila* peripheral nervous system (PNS) regenerate the same number of dendrites that an age-matched uninjured neuron has at that developmental time point.^14–16^ Dendrites can regenerate in the adult *Drosophila PNS*—even in aged adults.^17^ A recent study in mice also showed that neurons regenerate dendrites following initial degeneration after brachial plexus injury.^18^ Neurons can regenerate dendrites in the central nervous system (CNS) too. Dendrites of neurons in the zebrafish spinal cord regenerate after dendrite injury.^19^ Although mammalian CNS regeneration is not spontaneous, insulin treatment has been shown to promote dendrite regeneration after axotomy-induced dendritic retraction in mouse retinal ganglion cells.^20^ The capability for neurons to regenerate their dendrites provides a potential avenue to treat neurodegenerative disease pathology by activating regeneration in degenerating neurons.

To investigate dendrite regeneration in neurodegenerative conditions, we use the *Drosophila melanogaster* dendritic arborization (da) neurons as a model system.^21^ *Drosophila* have been important for identifying evolutionarily conserved genes and pathways in mammals and for modeling different diseases and have therefore proven an invaluable model to study neurodegenerative disease.^22–29^ Among the neurodegenerative diseases that have been modeled in *Drosophila*, polyglutamine (polyQ) diseases have been modeled extensively.

Polyglutamine diseases are a group of 9 known inherited autosomal dominant neurodegenerative disorders caused by expanded polyglutamine (CAG) repeats in a particular gene and include HD, SCAs, and spinal and bulbar muscular atrophy (SBMA). Short polyglutamine stretches, often 20 or fewer, occur naturally, but their expansion past a certain point, such as 30 or more, in genes related to these disorders leads to pathogenesis (though exact numbers vary based on the affected gene).^30^ In *Drosophila*, expression of human genes with expanded polyglutamine repeats recapitulates human disease pathology such as polyglutamine aggregates, neurodegeneration, dendrite defects, cytoskeletal aberrations, cell type-specific toxicity, behavioral defects, and RNA toxicity.^31–41^ In this paper we use *Drosophila* models for SCA1, SCA2, SCA3 and HD to study regeneration in the context of neurodegenerative disease.^31,34,40,42^

Here we show that degenerating neurons are capable of dendrite regeneration. We further show that injury to a single primary dendrite branch of neurons modeling neurodegenerative diseases induces a neuroprotective response in the rest of the dendrite arbor. Our work demonstrates that this effect is mechanistically tied to stabilization of the actin (but not microtubule) cytoskeleton. We show that this response is specific to dendrite injury, as axon injury and general injury to the surrounding tissue do not induce the same neuroprotection. Our results suggest that triggering dendrite regeneration can potentially act to ameliorate dendrite loss caused by neurodegeneration and further reveals key differences between dendrite and axon regeneration mechanisms.

## RESULTS

### Class IV da neurons overexpressing pathogenic polyglutamine transgenes experience progressive dendrite degeneration

We began by identifying neurodegenerative diseases where the dendrite degeneration can be readily studied. We can easily observe the dendrite arbors of the da neurons of the *Drosophila* PNS in intact animals. We first sought to characterize the effects of expressing different human polyglutamine (polyQ) transgenes in the highly branched class IV da neurons (ddaC) and the very simple class I da neurons (ddaE) of the *Drosophila* PNS in intact animals. In agreement with previous results,^35^ we found that expression of expanded polyglutamine disease transgenes MJD.78Q and ATX1.82Q in class 1 da neurons caused no obvious differences in dendrite arbors up to 120 hours after egg laying (AEL) (Figure S1G) but did cause dendrite defects in class IV da neurons (see below). We therefore decided to focus on class IV ddaC neurons for our study.

We expressed expanded polyQ transgenes modeling spinocerebellar ataxia type 1 (SCA1, ATX1.82Q), spinocerebellar ataxia type 2 (SCA2, ATX2.64Q), spinocerebellar ataxia type 3 (SCA3, MJD.78Q), and Huntington’s disease (HD, HTT231NT.128Q) in class IV da neurons. We compare these across time, from 24 hours AEL to 168 hours AEL, and to control neurons.

Wild type neurons add dendrite branches and dendrite length as well as increase in arbor complexity throughout larval development (Figure 1A,B,C and Figure S1A,C,D,E). We found that class IV ddaC neurons expressing any of the four long polyQ transgenes initially develop dendrites somewhat normally, but then later degenerate, losing dendrite branches and, in most cases, dendrite length. At 24 hours AEL, neurons overexpressing expanded polyglutamine disease transgenes were indistinguishable by eye from WT neurons and did not have significantly different dendrite branch number or length (Figure 1B and Figure S1A,F). Significant defects in dendrite arbor complexity measured by Sholl analysis in polyQ neurons compared to WT were first observed at 72 hours AEL and significant defects in counting branch number and measuring total dendrite length were first observed at 96 hours AEL (Figure 1B,C and Figure S1C,D,F). Not only are they different from wild type, but all models of polyglutamine disease exhibited the progressive loss of existing dendrites. In other words, we observe not just a slowing in dendrite addition, or just a stabilization of existing branches, but rather a decrease in total branch number over time. We quantified this true dendrite degeneration as a starting branch number at 24 hours AEL, a peak at some point (dendrite branch number peaks at 72-144 hours AEL, depending on the transgene), followed by a significant decrease in branch number by the last time point at 168 hours AEL (Figure S1E). All of our polyglutamine disease models also lost total dendrite length after a peak, except for ATX2.64Q. Expression of these constructs also caused degeneration of the axon terminals in the larval ventral nerve cord (Figure S1B).

**Figure 1.**
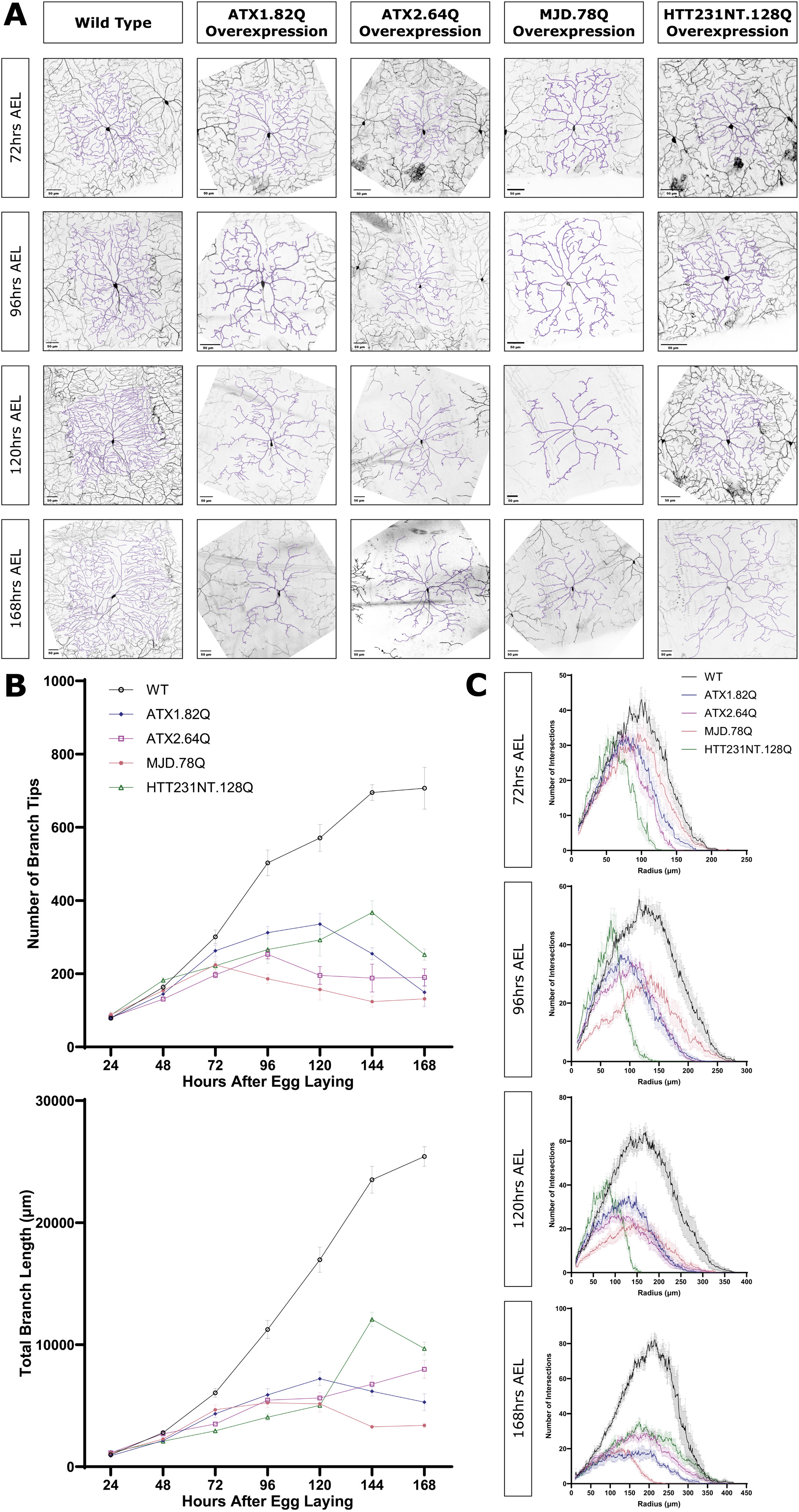
Overexpressing pathogenic polyQ proteins causes progressive degeneration of dendrites in class IV neurons. A) WT neurons and neurons overexpressing ATX1.82Q, ATX2.64Q, MJD.78Q, and HTT231NT.128Q at 72, 96, 120, and 168 hrs AEL. Scale bar 50 μm. B) Number of branches (top) and total branch length (bottom) at 24-168 hrs AEL for WT neurons and neurons overexpressing ATX1.82Q, ATX2.64Q, MJD.78Q, and HTT231NT.128Q. Mean ± SEM. C) Sholl analysis profiles at 72, 96, 120, and 168 hrs AEL for WT neurons and neurons overexpressing ATX1.82Q, ATX2.64Q, MJD.78Q, and HTT231NT.128Q. Mean ± SEM. Legends apply for all graphs in a panel. See also Figure S1.

While all four polyglutamine disease transgenes cause dendrite loss, there are some differences between the transgenes. MJD.78Q and ATX2.64Q cause dendrite loss at earlier stages (decreasing dendrite number beginning after 72-96 hours AEL), while ATX1.82Q and HTT231NT.128Q only begin to decrease dendrite number in older animals (decreasing dendrite number beginning after 120-144 hours AEL). Unlike the other three transgenes, ATX2.64Q overexpression does not cause shortening of dendrite length, though it does decrease dendrite number. Sholl analysis of dendrite arbor complexity seems to be the most sensitive method of quantification, detecting subtle defects at most or all time points, while dendrite branch number and dendrite length changes require a more obvious phenotype. Across all our quantification metrics, HTT231NT.128Q overexpression causes the mildest degeneration, MJD.78Q overexpression causes the most severe degeneration, and ATX1.82Q and ATX2.64Q overexpression each cause moderate degeneration.

Interestingly, although polyQ neurons were mostly indistinguishable from WT at 24 hours AEL, a transient increased arbor complexity was observed by Sholl analysis for ATX2.64Q and HTT231NT.128Q neurons that goes away by 48 hours AEL (Figure S1C,D). These results are consistent with emerging evidence that neurodegenerative polyglutamine diseases may have neurodevelopmental components including alterations in dendrite development.^43–49^

Overall, while WT ddaC neurons grow as larvae develop, class IV ddaC neurons overexpressing pathogenic polyglutamine proteins grow new dendrite branches and length until a point where they begin to degenerate dendrites and arbor complexity. Although the rate of degeneration differs between transgenes, these results indicate that polyglutamine model neurons allow us to study how neurodegenerative dendrite loss is affected by regeneration following dendrite injury.

### Neurons overexpressing pathogenic polyglutamine proteins can regenerate dendrites following severe injury

Previous work in da neurons demonstrated that class IV ddaC neurons can regenerate branch number but not branch length within 72 hours after severe injury when all dendrites are severed (balding) at 48 hours after egg laying.^14^ We investigated whether neurons overexpressing polyglutamine disease transgenes were capable of regenerating dendrites after injury. To do this we balded neurons at 48 hours AEL, tracked regeneration from 24 to 72 hours after injury (AI), and compared to age-matched uninjured control neurons.

As expected, WT neurons that were balded at 48 hours AEL were able to regenerate branch number but not branch length between 24 and 72 hours AI (Figure 2B,C,D). In the absence of injury, MJD.78Q neurons lose branches and uninjured ATX1.82Q neurons stagnate during this same time (Figure 2A,C). Surprisingly, we found that overexpression of ATX1.82Q and MJD.78Q did not preclude neurons from regenerating dendrite branches after balding during the same period, and balded ATX1.82Q and MJD.78Q neurons added branch number and length (Figure 2A,C). The quality of regeneration depended on the transgene. Neurons overexpressing MJD.78Q could regenerate both dendrite branch number and total branch length enough to match uninjured age matched control neurons by 72 hours AI (Figure 2D). Neurons overexpressing ATX1.82Q were able to regenerate, but not enough to match uninjured age matched controls in branch number or length (Figure 2D). We also observed regeneration after balding at 48 hours AEL for HTT231NT.128Q neurons (Figure S2A). Following full balding at 96 hours AEL, ATX1.82Q neurons were capable of mild regeneration though MJD.78Q neurons were not (Figure S2B). This result demonstrates that polyQ model neurons are capable of regenerating dendrites following full balding injury at 48 hours AEL and that degeneration does not completely prevent dendrites from regenerating at 96 hours AEL. These results suggest that dendrite *de*generation caused by polyglutamine disease transgene expression does not inhibit dendrite *re*generation pathways.

**Figure 2.**
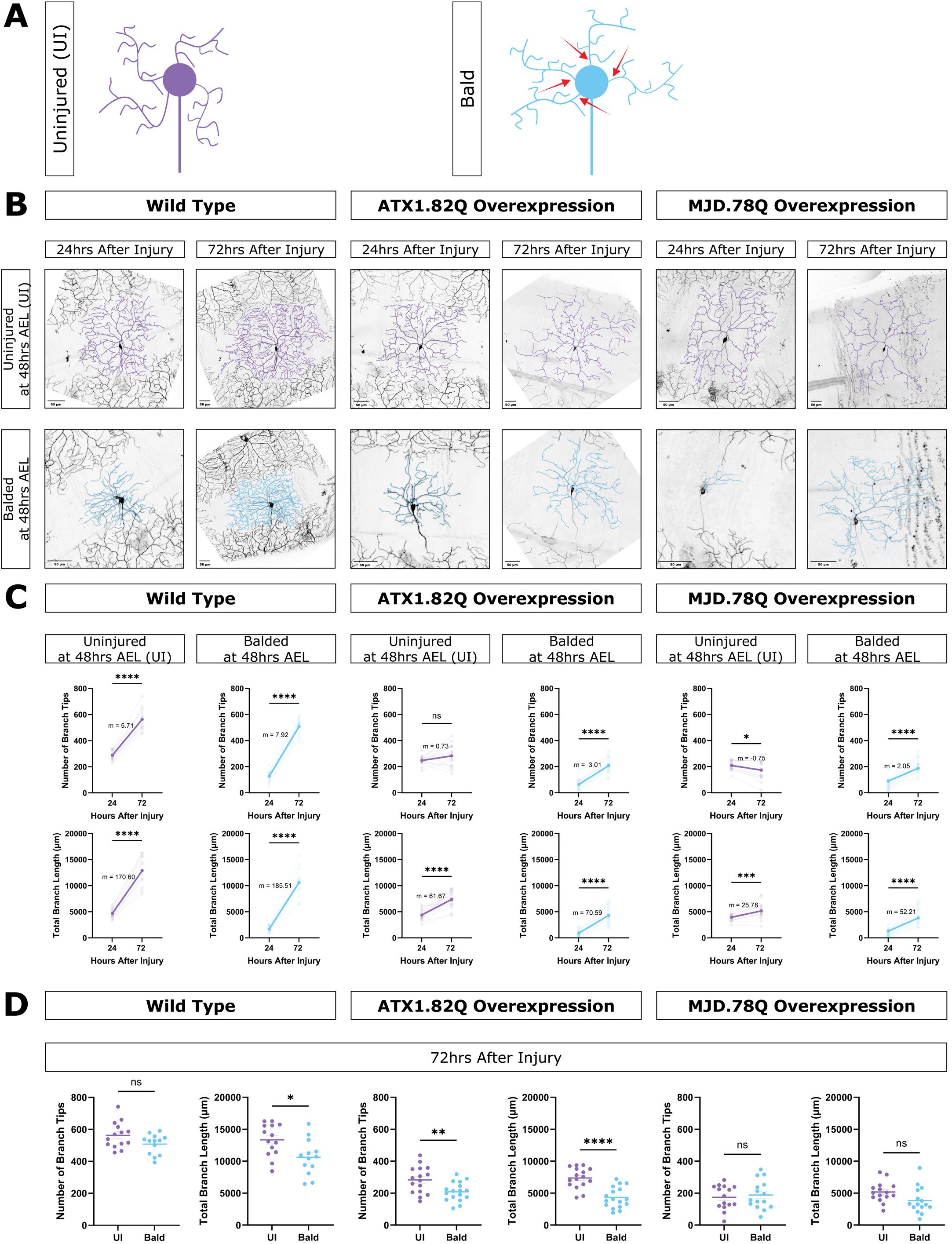
Neurons overexpressing pathogenic polyQ proteins can regenerate dendrite arbors following complete removal at 48 hours AEL. A) Schematic of uninjured neurons (purple) and bald neurons (blue). B) Uninjured neurons and balded neurons for WT, ATX1.82 overexpression, and MJD.78Q overexpression neurons at 24 and 72 hrs after injury done at 48 hrs AEL. Scale bar 50 μm. C) Number of branch tips (top) and total branch length (bottom) at 24 and 72 hrs after injury for uninjured (purple) and balded (blue) for WT, ATX1.82Q, and MJD.78Q overexpression neurons. Individual neurons (faded) and mean (bold) values shown, with slope (m) between the mean 24 and 72 hour AI values. Paired t-test. D) Number of branch tips (left) and total branch length (right) between uninjured and balded WT, ATX1.82Q, and MJD.78Q overexpression neurons at 72 hrs AI. Welch’s t-test. See also Figure S2.

### Injury to a single dendrite branch induces neuroprotection in neurons overexpressing pathogenic polyglutamine proteins

Having established that neurons overexpressing expanded polyglutamine disease transgenes are capable of regenerating dendrites, we sought to understand if this regenerative capacity can rescue degeneration at later developmental time points. To assess this, we injured neurons at a time point when polyQ model neurons had significantly fewer dendrite branches and shorter length compared to WT, around 96 hours AEL, which marked the start of the degenerative phase of neuron growth (as demonstrated previously, in Fig 1 and Supp Fig 1). To determine if dendrite regeneration could alter this degenerative fate, we performed single primary dendrite branch injuries (SBI) to neurons and compared them to age-matched normalized control neurons (NC). Normalized controls were quantified to normalize for the loss of one primary branch by not including that primary branch and its higher order branches in the quantification (Figure 3A).

**Figure 3.**
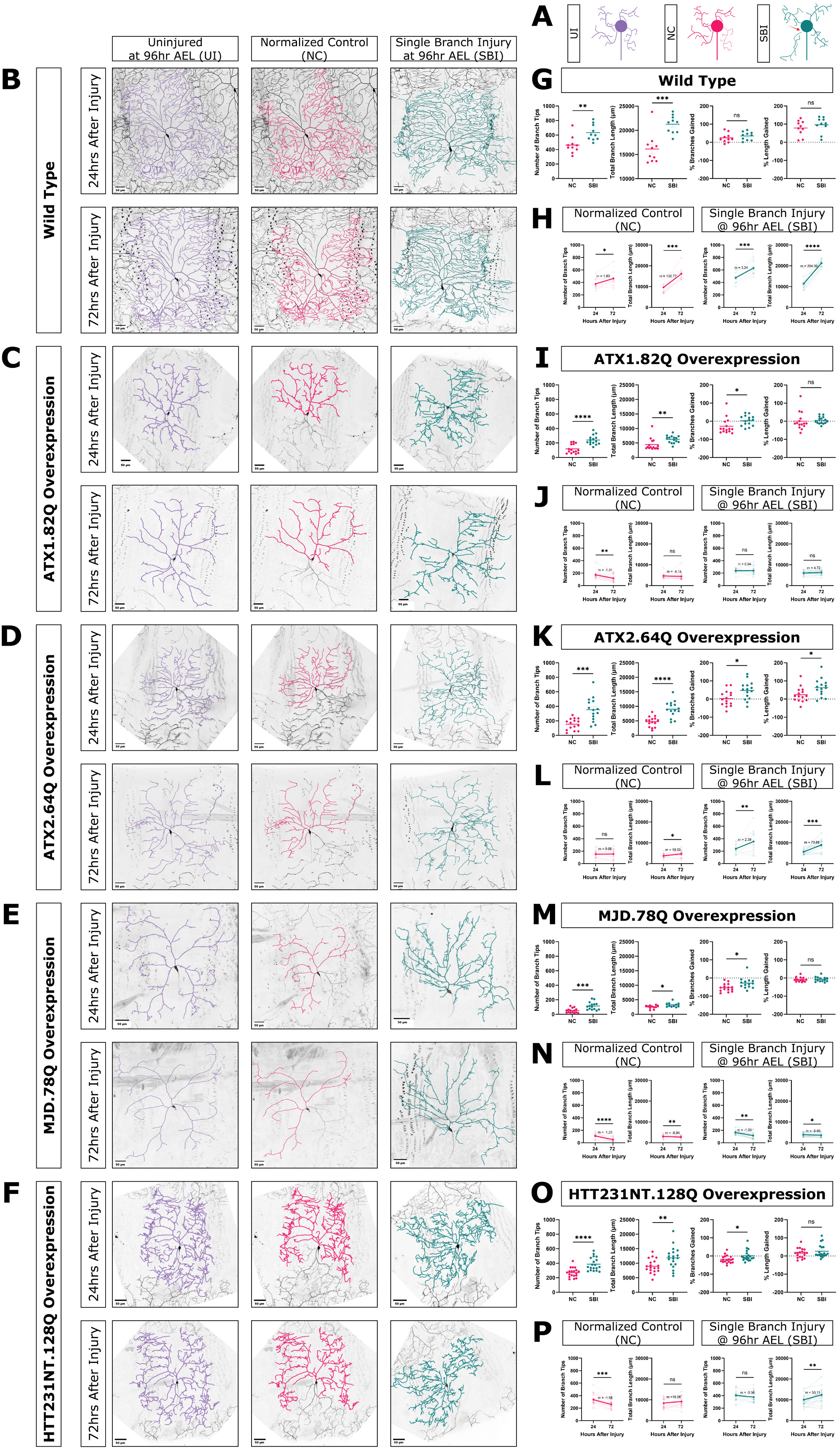
Injury to a single dendrite branch induces neuroprotection in neurons overexpressing pathogenic polyQ proteins. A) Schematic of uninjured neurons (purple, UI), normalized control neurons (pink, NC), and single branch injured neurons (green, SBI). B-F) Uninjured neurons, normalized control neurons, and single branch injured neurons at 24 and 72 hrs after injury for WT, ATX1.82Q, ATX2.64Q, MJD.78Q, and HTT23NT.128Q overexpression neurons. Scale bar 50 μm. G, I, K, M, O) Number of branch tips and total branch length of NC and SBI neurons at 72 hrs after injury, or comparing % change in number of branch tips and total branch length between 24 to 72 hrs after injury between NC and SBI neurons. Welch’s t-test. H, J, L, N, P) Number of branch tips and total branch length at 24 and 72 hrs after injury for NC (pink) and SBI (green) neurons. Individual neurons are faded, solid lines represent the average slope (m) between the mean 24 and 72 hour AI values. Paired t-test. See also Figure S3.

We found that WT SBI neurons had significantly more dendrite branches and branch length than NC neurons at 72 hours AI (Figure 3G). Both WT SBI neurons and WT NC neurons had significant increases in branch number and length between 24 and 72 hours AI (Figure 3H). When we determined the amount of branches or length added between 24 to 72 hours AI compared to the starting number of branches at 24 hours AI, there was no significant difference in either percent branches added or percent length added between WT SBI and NC neurons, indicating that growth in SBI neurons was proportional to the number of branches and length the neurons started with after injury (Figure 3G). In other words, the growth in WT SBI neurons was equivalent to developmental growth of age-matched control neurons during the same time.

Next, we examined the response of polyQ neurons to single branch injury, compared to normalized controls. We first looked at the terminal time point, 72 hours AI. Like WT SBI neurons, all polyQ model SBI neurons had significantly more dendrite branches and length at 72 hours AI than NC neurons (Fig 3I,K,M,O). We next looked at growth and degeneration during the time between injury and the terminal time point. Unlike WT SBI neurons, all polyQ model SBI neurons added more branches proportional to their starting branches than polyQ NC neurons (Fig 3I,K,M,O). This trend was also true for added length in the case of ATX2.64Q neurons (Fig 3K). This indicates that, unlike in WT neurons, the growth experienced by polyQ SBI neurons was different from the stagnation or degeneration seen in age-matched control neurons. Additionally, we observed that location of the injured branch did not affect the location of new growth in the rest of the arbor (Figure S3F,G).

The effect of single branch injury on the rate of branch number change and length change during this time differed depending on the transgene. For ATX1.82Q and HTT231NT.128Q neurons, we observed that single branch injury rescued the significant decrease in branch number seen in their paired NC neurons, inducing branch number stagnation rather than branch loss during that time (Figure 3J,P). For ATX2.64Q neurons, single branch injury induced a significant increase in branch number compared to stagnation for NC neurons (Figure 3L). The effect of SBI on MJD.78Q neurons was weaker, with both SBI and NC neurons decreasing in both branch number and length, though at a slower rate for the SBI neurons (Figure 3N). Overall, we observed that single branch injury was able to alter, and in some cases rescue, the dendrite loss trajectory of pathogenic polyQ neurons between 24 and 72 hours AI.

To determine whether this was specific to degenerating polyQ neurons, we also examined responses to SBI in two non-pathogenic polyQ overexpression models, ATX1.30Q and MJD.27Q. We found that ATX1.30Q appeared like WT in the uninjured condition and in response to SBI (Figure S3C). We also found that MJD.27Q neurons exhibited variable degeneration at late time points, but responded to SBI similarly to WT neurons (Figure S3A,B). We also examined response to SBI in a non-polyQ-induced model of dendrite degeneration by overexpressing a transgene for the 0N3R isoform of human tau (MAPT)^50^ in class IV neurons and conducting single branch injuries at 96 hours AEL. We found that single branch injured hMAPT.0N3R overexpression neurons also had significantly more dendrite branches than normalized controls (Figure S3D,E), suggesting that the single branch injury-induced neuroprotection may be applicable in other models of neurodegenerative disease.

Altogether, we observed that single branch injury induces regenerative growth that increased dendrite number and length in pathogenic polyQ model neurons. Although there were differences between individual transgenes, single branch injury activated regeneration across all polyQ models we tested, leading to protection of existing branches or growth of new branches. This supports the idea that injury of one dendrite branch can effectively “turn on” regenerative processes in uninjured branches, to slow or limit their degeneration. We refer to this phenomenon as single branch injury induced neuroprotection.

### Axotomy does not induce neuroprotection in neurons overexpressing pathogenic polyglutamine proteins

We next tested if the neuroprotection was specific to dendrite injury by instead injuring axons. To assess this, we performed axotomies at ∼40µm from the cell body at 96 hours AEL and assessed dendrite architecture at 24 and 72 hours AI. We performed axotomies at this distance to avoid dendrite to axon transition during axon regeneration.^51–53^ WT neurons lost dendrite branches and length following axotomy, and axotomized neurons had significantly fewer dendrite branches and length than age-matched uninjured control neurons at 24 hours AI (Fig 4B,C,D). This is in agreement with previous observations in our system and other species.^54–58^ Axotomized WT neurons lost branches and length between 24 and 72 hours AI while uninjured age-matched control WT neurons grew branches and length during that time (Fig 4C).

**Figure 4.**
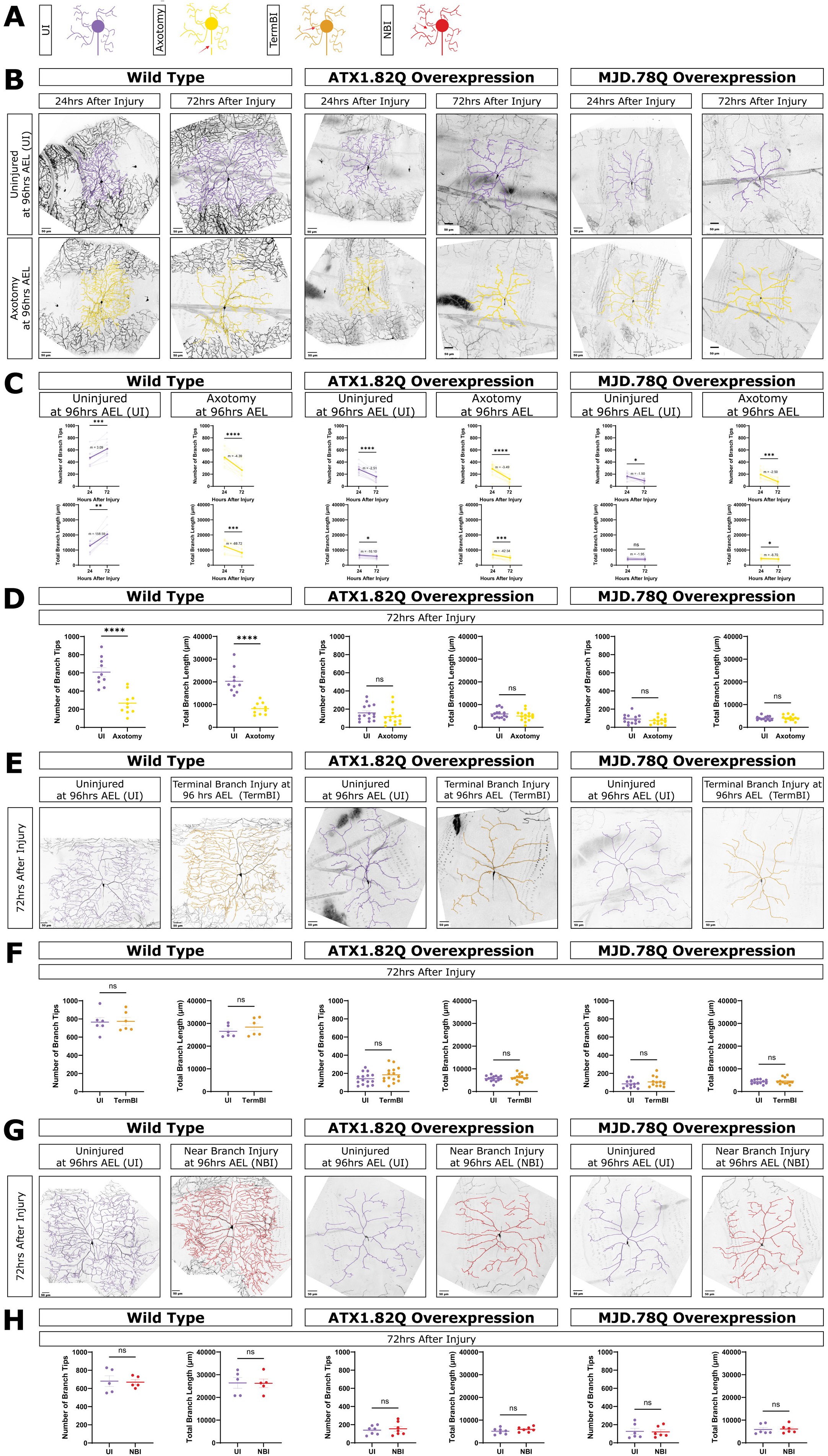
Axotomy, terminal branch injury, and near branch injury do not induce neuroprotection. A) Schematic of uninjured neurons (purple, UI), axotomized neurons (yellow), terminal branch injured neurons (orange, TermBI), and neurons left uninjured but exposed to an injury near a dendrite branch (red, NBI). B) Uninjured and axotomized WT, ATX1.82Q, and MJD.78Q overexpression neurons at 24 and 72 hrs after injury. Scale bar 50 μm. C) Number of branch tips (top) and total branch length (bottom) at 24 and 72 hrs after injury for uninjured (purple) and axotomized (yellow) neurons. Paired t-test. D) Number of branch tips and total branch length between UI and axotomy at 72 hrs AI. Welch’s t-test. E) Uninjured and terminal branch injured neurons at 72 hrs AI. Scale bar 50 μm. F) Number of branch tips and total branch length between UI and TermBI at 72 hrs AI. Welch’s t-test. G) Uninjured and uninjured but exposed to a near branch injury neurons at 72 hrs AI. Scale bar 50 μm. H) Number of branch tips and total branch length between UI and NBI at 72 hrs AI. Welch’s t-test.

We found that axotomy of neurons overexpressing ATX1.82Q and MJD.78Q did not produce any neuroprotection. Axotomized ATX1.82Q neurons and axotomized MJD.78Q neurons both lost branches and length between 24 and 72 hours AI, similar to uninjured neurons expressing these pathogenic transgenes (Fig 4C,D). This shows that axon injury does not activate a neuroprotective effect in degenerating polyQ neurons, unlike what was observed with dendrite branch injury, establishing the neuroprotective effect as specific to dendrite injury.

### Injury-induced neuroprotection is specific to primary branch injury and is cell autonomous

Having established that only dendrite injury activates a neuroprotective effect, we also tested whether a small injury to a terminal dendrite branch could recapitulate the effect of a single primary dendrite branch injury. For WT neurons, we did not observe a significant difference in branch number or length between uninjured age-matched control neurons and terminal branch injured neurons at 72 hours AI (Fig 4E,F). Similarly, for both ATX1.82Q and MJD.78Q, terminal branch injury was not sufficient to induce the neuroprotective effect seen with single primary branch injury (Fig 4E,F). Overall, terminal branch injury was not sufficient to induce growth for WT neurons or neuroprotection for polyQ neurons.

To test if the observed neuroprotective effect was cell autonomous, or perhaps a result of injuring surrounding cells such as epidermal cells, we also conducted experiments where the laser was aimed at a space near a primary dendrite branch at a similar distance from the cell body as SBI experiments. In WT neurons, neurons that were exposed to a near-miss injury were not significantly different in branch number or length at 72 hours AI from age-matched uninjured neurons with no laser injury at all (Fig 4G,H). Similarly, a near-miss injury did not produce a similar neuroprotective effect in either ATX1.82Q or MJD.78Q neurons (Fig 4G, H). This data suggests that the single primary dendrite injury induced neuroprotection we observed is due to processes being initiated inside the injured cell, leading to growth and retention of branches, rather than a result of surrounding cells.

### Single branch injury, but not axotomy, induces stabilization of the actin, but not the microtubule, cytoskeleton

We next wanted to determine a mechanistic explanation for the SBI induced regenerative neuroprotection we observed in polyQ neurons, so we looked at microtubules (MT) and F-actin in these neurons after injury. We hypothesized that single branch injury-induced regeneration might be triggering stabilization or rescue of actin and MTs in degenerating polyQ neurons, leading to neuroprotection.

First, we looked at actin in the neurons. To look at F-actin in these neurons, we chose to use the construct GMA, a fusion protein which consists of GFP fused to the cytoskeletal linker protein moesin’s actin-binding domain, that directly labels F-actin during nucleation.^59–61^ This construct shows areas with stable actin and was expressed only in the class IV da neurons. To determine if F-actin stabilization might play a role in the single branch injury-induced neuroprotective response that we observed, we injured neurons and quantified the GMA signal in the cell body and dendrite branches at 24 hours AI.

In WT neurons, we found that single branch injury did not change stable F-actin in either the cell body or dendrite branches, and that axotomy significantly reduced F-actin levels in both the cell body and dendrite branches (Figure 5A,B). In contrast, for ATX1.82Q, ATX2.64Q, and HTT231NT.128Q neurons, single branch injury caused an increase in F-actin in the cell body and dendrite branches (Figure 5A,B). Axotomy however reduced F-actin in the cell body of ATX2.64Q and HTT231NT.128Q neurons and did not change F-actin in ATX1.82Q neurons (Figure 5A,B). Axotomy did not change F-actin in branches in ATX1.82Q, ATX2.64Q, and HTT231NT.128Q neurons (Figure 5A,B). These results show that single branch injury triggers stabilization of F-actin in both dendrite branches and in the cell body for degenerating neurons, while axotomy does not. Our results show that axon injury leads to loss of F-actin in the cell body and dendrites whereas dendrite injury leads to an increase in F-actin in both the cell body and dendrites, but this is only seen in neurons already defective for F-actin, such as polyQ model neurons. This data suggests that single branch injury causes regenerative neuroprotection in degenerating neurons by leading to stabilization of the actin cytoskeleton and that this does not occur following axotomy.

**Figure 5.**
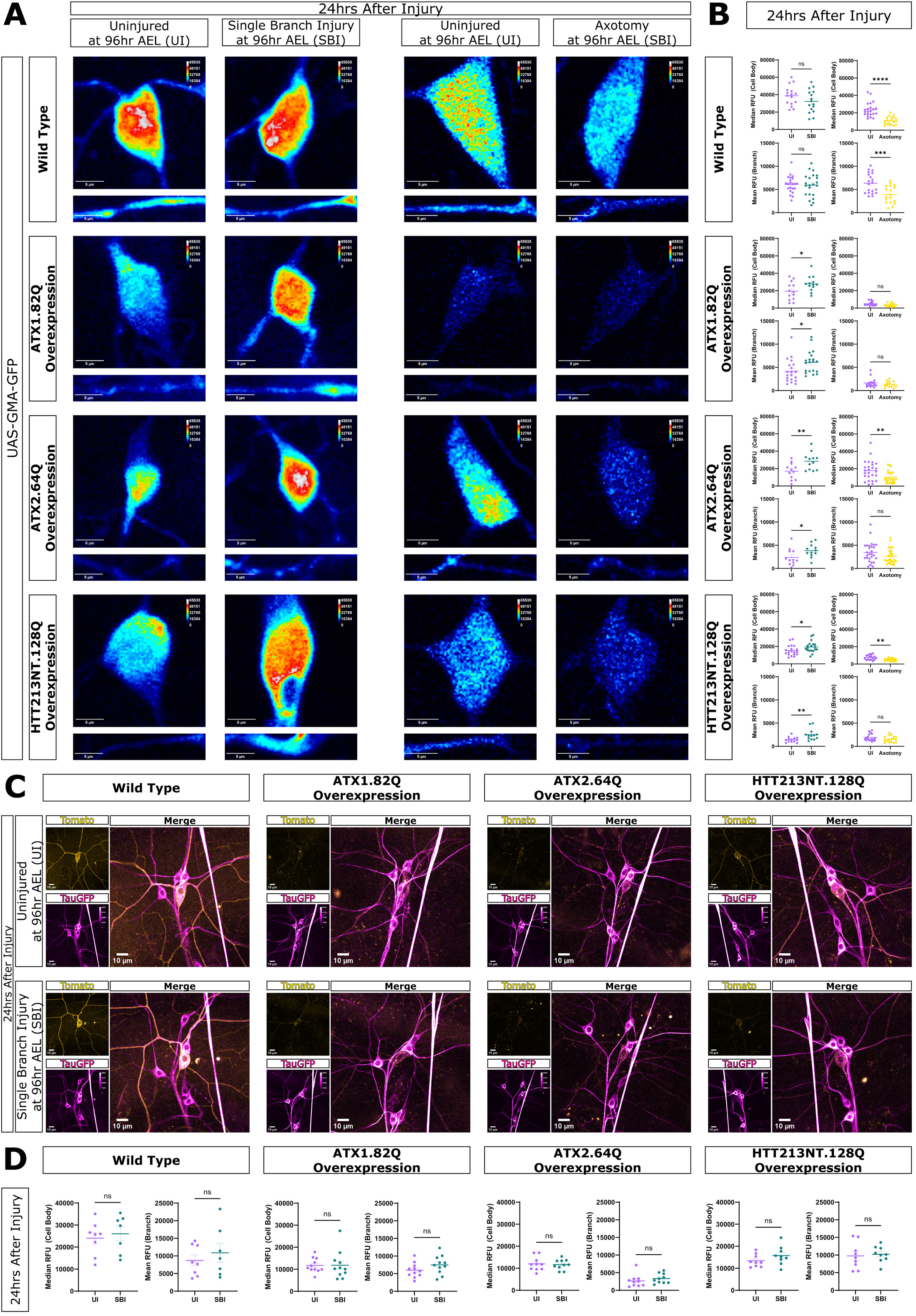
Single branch injury, but not axotomy, induces stabilization of the actin, but not the microtubule, cytoskeleton. A) GMA imaging at 24 hrs after injury for WT, ATX1.82Q, ATX2.64Q, and HTT231NT.128Q neurons. Images (L>R): uninjured control cell body (top) and branch (bottom) images for SBI experiments, SBI cell body and branch, uninjured control for axotomy experiments, axotomy cell body and branch. Scale bar 10 μm. B) Median cell body RFU (top) and mean branch RFU (bottom) between uninjured versus SBI neurons (left) and between uninjured versus axotomy neurons (right) for WT, ATX1.82Q, ATX2.64Q, and HTT231NT.128Q neurons. Welch’s t-test. C) TauGFP imaging at 24 hrs after injury for uninjured, SBI, and axotomy of WT, ATX1.82Q, ATX2.64Q, and HTT231NT.128Q neurons. Ppk-CD4-tdtomato pseudo-colored in yellow, Tau-GFP pseudo-colored in magenta. D) Median cell body RFU and mean branch RFU between uninjured versus SBI neurons for WT, ATX1.82Q, ATX2.64Q, and HTT231NT.128Q neurons. Welch’s t-test.

Next, we wanted to determine if the neuroprotective mechanism was specific to actin stabilization or if it also involved changes in MT stability and dynamics. We examined microtubules using three different approaches: visualizing Tau-GFP, immunostaining for futsch/MAP1b, and observing new MT growth using EB1-GFP comets. We first looked at stable microtubule bundles using the construct Tau-GFP, which labels endogenous microtubules in both axons and dendrites of all da neurons.^62–64^ We injured and subsequently imaged neurons with ppk-CD4-tdTomato and used Tau-GFP to visualize microtubules 24 hours AI. We found that single branch injury did not change the amount of Tau-GFP in either dendrites or the cell body for WT or polyQ model neurons (Figure 5C,D). We next immunostained for *Drosophila* microtubule associated protein futsch/MAP1b at 24 hours AI and did not see an effect on microtubule localization or density (Supp Figure 5E). Finally, we examined microtubule dynamics by imaging EB-1 comets in neurons 24 hours AI. Interestingly, single branch injury induced mixed microtubule polarity in both WT and polyQ neurons (Supp Fig 5D). In addition, single branch injury rescued the significantly increased microtubule dynamics caused by MJD.78Q overexpression (Supp Fig 5C). However, the same was not observed for ATX1.82Q neurons where single branch injury did not affect comet number or velocity (Supp Fig 5B). This indicated that, although single branch injury did induce some changes in microtubule dynamics, new microtubule formation or microtubule stabilization likely do not contribute to a core mechanism behind the observed neuroprotection.

Together these data suggest that the mechanism of single branch injury-induced neuroprotection is likely stabilization of F-actin. Our observations also suggest that actin stabilization occurs early in the regenerative process after injury and this early stabilization is enough to stabilize the arbor for sustained neuroprotection we observed later at 72 hours AI.

## DISCUSSION

Our study presents the first evidence that dendrite regeneration is possible in degenerating neurons and demonstrates that dendrite regeneration triggered by single branch injury can induce neuroprotection of the remaining dendrite arbor in ddaC neurons modeling neurodegenerative diseases. We demonstrate that this effect is cell autonomous and specific to primary dendrite branch injury—not axon injury. Our results also show that this neuroprotection is supported by the mechanism of early stabilization of the actin cytoskeleton in both dendrite branches and the cell body.

### Neurodegeneration does not prevent regeneration

Our work demonstrates that although neurodegeneration leads to progressive dendrite loss, it does not prevent dendrite regeneration following injury. We demonstrate that both neurons destined to degenerate and neurons actively degenerating are capable of some regeneration, with varying success depending on the type of dendrite injury, the stage of degeneration, and the disease model. We further show that this effect not only applies to polyglutamine disease models, but another neurodegenerative model caused by overexpression of human tau. Together, this data suggests that neurons demonstrate capacity to recover from neurodegeneration, that neurodegeneration does not prevent regeneration, and that dendrite degeneration in neurodegenerative disease is reversible.

### The cytoskeleton in regeneration and neurodegenerative disease

Proper MT and F-actin regulation are critical for neurite growth, morphology and stability.^35,65^ Enrichment of local F-actin is associated with dendritic branching, and dendritic arbor length is highly interrelated with local MT quality.^65^ In addition, the importance of actin nucleator Cobl in mice and the actin regulators RAC GTPase CED-10 and RhoGEF TIAM-1 in *C. elegans* for dendrite regeneration suggests a key role for the actin cytoskeleton in the regeneration process of dendrites.^66,67^ Similarly, the MT binding protein Patronin and its microtubule nucleation function have been implicated in dendrite regeneration.^68–70^ Our results suggest that stabilization of F-actin plays an important role in the response to dendrite injury and subsequent regeneration. Although single branch injury had variable effects on stable microtubules and dynamics, we found that dendrite injury induced mixed microtubule polarity in all neurons, suggesting that changes in microtubule polarity play a role in response to dendrite injury and dendrite regeneration.

In accordance with the importance of the cytoskeleton for neuron development and stability, cytoskeletal dysregulation is a hallmark of neurodegenerative disease.^71–77^ Regulators of the cytoskeleton such as cofilin, RhoA/ROCK pathway components, MAPs, GSK3β have emerged as potential therapeutic targets for treating this dysregulation.^74,77–81^ It has previously been shown that F-actin levels are reduced in SCA1 and SCA3 model class IV da neurons.^35^ Our results suggest that this reduction can be ameliorated by activating dendrite regeneration mechanisms via single branch injury. This indicates that dendrite regeneration mechanisms present another pathway that may be targeted to modulate the actin cytoskeleton for therapeutic potential in neurodegenerative disease.

### Insights about dendrite and axon regeneration

Our work demonstrates a unique phenomenon that can be triggered by dendrite injury and dendrite regeneration mechanisms, but not for axon injury and axon regeneration mechanisms, highlighting the need for more focus on the molecular underpinnings of dendrite regeneration. Much focus has been placed on studying axon injury and regeneration, with little attention paid to dendrite injury and regeneration, and questions remain about how much these two processes share or diverge. Although dendrite and axon regeneration after injury share certain mechanistic aspects, such as regulation by the Akt pathway, other aspects have emerged as unique to one process and not the other.^16^ For example, axon regeneration is triggered via a conserved signaling cascade that relies on the dual leucine zipper kinase DLK, but DLK and its downstream transcription factors have also been shown to be dispensable for dendrite regeneration.^15,82–87^ Similarly, dendrite regeneration in *Drosophila* requires the tyrosine kinase Ror, but this kinase is not required for axon regeneration.^70^ Our study adds to this body of evidence by showing that dendrite regeneration mechanisms, and not axon regeneration mechanisms, have unique potential to protect dendrite arbors in degenerating neurons. Our work also demonstrates a notable mechanistic difference between axon and dendrite regeneration. In axotomized neurons, F-actin in the cell body and dendrite arbor is reduced early in the regeneration time frame, while the opposite seems to be triggered following dendrite injury. Although we did not see an increase in F-actin in WT neurons following single branch injury, our results in polyQ model neurons suggests that this increase may only be observable in neurons where F-actin levels are already lowered or dysregulated. Our work along with others demonstrates a growing need for deeper investigation into mechanisms behind dendrite regeneration to both inform our understanding of regenerative processes that may be harnessed in neurons and to elucidate new avenues of exploration in diseases that affect dendrites.

### Is dendrite regeneration just upregulation of dendrite maintenance mechanisms?

Another question that remains in the field of dendrite regeneration is whether dendrite regeneration is entirely a result of upregulated dendrite maintenance mechanisms, has some overlap with dendrite maintenance mechanisms, or is its own completely separable process. Although it is known that this regeneration is different from developmental growth of dendrites, it is not understood how much these processes of development versus regeneration overlap and whether something that prevents proper development and maintenance, such as neurodegenerative disease, will also prevent regeneration. Both how dendrites are maintained throughout life and how they can regenerate after injury is largely understudied and little is known about how these two processes may be either interconnected or distinct from one another.

Growing dendrite arbors are dynamic and their continual extension and retraction until they reach maturity is influenced by synaptic connections.^88–90^ After development, dendrites and synapses must be actively maintained with long-term stability regulated by a combination of signaling pathways, synaptic inputs, and structural support from the cytoskeleton and cell adhesion and scaffolding molecules.^88–90^ Our work suggests that dendrite maintenance and dendrite regeneration are indeed separable processes because defective maintenance does not preclude regeneration or regrowth after injury. We demonstrate that neurons that fail to maintain their dendrite arbors due to neurodegenerative dendrite loss are still capable of regeneration. Although the regeneration process after severe balding injury is stunted in degenerating neurons at later stages of degeneration, we still observed some regrowth. Because defective maintenance in our study is due to neurodegeneration that perturbs many aspects of neuronal homeostasis, health, and maintenance, further exploration looking at dendrite regeneration in neurons where dendrite maintenance mechanisms are more directly inhibited is needed to distinguish the two processes more definitively.

### Beneficial outcomes of stress

Moderate stressors have been associated with benefits for organisms. Previous work by Neumann et. al demonstrated that introducing priming lesions both at the time of spinal cord injury and a week later were able to enhance axonal regeneration in adult rat DRG neurons.^91^ Acute stress has also been shown to increase neurogenesis in the adult rat hippocampus, leading to benefits in learning and memory tests.^92^ Mild stress due to dietary restriction has shown benefits for lifespan, health span and brain health.^93–95^ Our work demonstrates a similar phenomenon in the context of neurodegenerative disease and dendrite repair. Previous work has shown that activating the integrated stress response is beneficial in HD and ALS models.^96–99^ In our study, we show that the stress of a subtle injury to a single dendrite can cause a beneficial regenerative response in the remaining arbor. Our work adds to the growing body of evidence that acute injury or transient stress can have beneficial outcomes for neuronal repair and health.

### Implications for and potential therapeutic avenues in neurodegenerative disease

Our results reveal an unexplored pathway for potential therapeutic targets in neurodegenerative disease. Our new findings provide the first evidence for potential regeneration of dendrites lost in neurodegenerative disease. Our work suggests that dendrite regeneration mechanisms can be harnessed to recover and preserve dendrites, potentially delaying or slowing the cellular effects of neurodegenerative disease and ultimately improving neuronal function. Further work to characterize the intermediate factors controlling the neuroprotective response that lead to downstream actin stabilization will be key to discovering new therapeutic targets in neurodegenerative disease. Perhaps if we can modulate these factors similarly but in the absence of direct neuronal injury, we can achieve sustained neuroprotection.

Our study also hints that just having the gain of function effect of expanded polyglutamine aggregate overexpression is not only enough to recapitulate neurodegenerative phenotypes, but also transient neurodevelopmental phenotypes. A growing amount of evidence suggests altered neurodevelopment due to loss of function of important genes for brain development like huntingtin is an important feature of disease pathogenesis in polyglutamine diseases and other neurodegenerative disorders.^43–49,100–102^ Our study suggests that, in addition to LOF effects, GOF effects may also play a role in neurodevelopmental defects observed in polyglutamine disease.

### Conclusions

Taken together, the results of our study support the idea that injury of one dendrite branch can effectively “turn on” regenerative processes in uninjured branches, leading to retention or new growth and slowing or limiting degeneration in spared branches. These results present a promising avenue to explore triggering dendrite regeneration as a possible treatment for degenerative dendrite loss in neurodegenerative diseases. Future work should focus on assessing these phenomena in adult mature neurons and in neurons of the CNS. Further studies have the potential to elucidate both important mechanisms underlying dendrite regeneration and potential therapeutic targets for intervention in neurodegenerative disease.

## ACKNOWLEDGEMENTS

Research in this publication was supported by NIH grant R00NS097627 (to KTP). Research reported in this publication was also supported by the California Institute for Regenerative Medicine under Award Number EDUC4-12822. The content is solely the responsibility of the authors and does not necessarily represent the official views of the California Institute for Regenerative Medicine. The authors would also like to acknowledge the University of California Irvine’s Undergraduate Research Opportunities Program and Summer Undergraduate Research Program (UROP and SURP). We thank Vicky Lam for other informative data analysis work not included here. We thank Vinicius Duarte, Rosty Brichko, and Mia Brantley for their intellectual contributions to this work. We thank Nancy Bonini, Juan Botas, and Melissa Rolls for providing certain *Drosophila* stocks. We thank the UCI Optical Biology Core and Dr. Adeela Syed for extensive use of the microscopes in their facility. This study was made possible in part through access to the Optical Biology Core Facility of the Developmental Biology Center, a shared resource supported by the Cancer Center Support Grant (CA-62203) and Center for Complex Biological Systems Support Grant (GM-076516) at the University of California, Irvine.

## AUTHOR CONTRIBUTIONS

Conceptualization, S.E.P. and K.T.P.; Methodology, S.E.P. and K.T.P.; Formal Analysis, S.E.P., I.N.B., D.S., G.E.J., A.M., L.U.D, H.H.N.L.; Investigation, S.E.P., I.N.B., D.S., P.T.H., J.G.M.; Resources, K.T.P.; Data Curation, S.E.P.; Writing – Original Draft, S.E.P.; Writing – Review & Editing, S.E.P., P.T.H., K.T.P.; Visualization, S.E.P.; Supervision, K.T.P.; Project Administration, S.E.P., K.T.P.; Funding Acquisition, S.E.P., K.T.P., I.N.B., D.S., A.M., L.U.D.

## STAR METHODS

### RESOURCE AVAILABILITY

#### Lead Contact

Further information and requests for resources should be directed to and will be fulfilled by the lead contact, Katherine Thompson-Peer.

#### Materials availability

No new unique reagents were generated by this study.

#### Data and code availability

All data reported in this paper will be shared by the lead contact upon request. This paper does not report original code. Any additional information required to reanalyze the data reported in this paper is available from the lead contact upon request.

### EXPERIMENTAL MODEL AND SUBJECT DETAILS

#### Drosophila strains

*Drosophila* stocks were maintained at room temperature. Larva for experiments were maintained at room temperature or in an incubator at 22.5°C and 70% humidity. Both male and female larva were used for all experiments. The following fly strains were used in this study: Canton-S, ppk-CD4-tdGFP (chromosome 3),^103^ ppk-gal4 (chromosome 2),^104^ ppk-cd4-tdTomato (chromosome 2),^103^ UAS-CD4-tdTomato (chromosome 2),^103^ Gal4^2-21 (chromosome 3),^105^ UAS-GMA (BDSC#31776), UAS-EB1-GFP (BDSC#35512), WeeP304(tau-GFP) (courtesy of Melissa Rolls),^63^ UAS-Hsap\HTT231NT.128Q (courtesy of Juan Botas),^42^ UAS-hATXN3.tr-Q78 (BDSC#8150), UAS-Hsap\ATX1.82Q (BDSC#33818), UAS-ATXN2-CAG-64 (courtesy of Nancy Bonini),^40^ UAS-hATXN3.tr-Q27 (BDSC#8149), UAS-hSap\ATX1.30Q (BDSC#39739), UAS-hMAPT.03NR (BDSC #93609). A complete list of fly stocks can be found in the key resources table.

### METHOD DETAILS

#### Generation of fly lines and experimental crosses

Crosses were performed at room temperature or in an incubator at 22.5°C and 70% humidity on a plate made of grape juice and agarose (grape plate) with yeast paste to synchronize animal age. Cross progeny larva for experiments were kept on grape plates at room temperature or in an incubator at 22.5°C and 70% humidity. Assays to assess development, injury, and regeneration in class IV ddaC neurons were performed by crossing fly lines expressing ppk-cd4-tdGFP and ppk-gal4 with CantonS flies or flies expressing UAS-polyQ. Assays to assess development in class 1 ddaE neurons were performed by crossing fly lines expressing Gal4^2-21 and UAS-tdTomato with CantonS flies or flies expressing UAS-polyQ. Assays to assess F-actin in class IV ddaC neurons were performed by crossing fly lines expressing UAS-GMA under control of ppk-gal4 with CantonS (WT) flies or flies expressing UAS-polyQ. Assays to assess microtubule dynamics in class IV ddaC neurons were performed by crossing fly lines expressing UAS-EB1-GFP under control of ppk-gal4 with CantonS flies or flies expressing UAS-polyQ. Assays to observe stable microtubule bundles in class IV ddaC neurons were performed by crossing fly lines expressing ppk-gal4, ppk-cd4-tdtomato and WeeP.304TauGFP with CantonS flies or flies expressing UAS-polyQ. Following injury and subsequent imaging, larva were individually housed on grape plates with yeast paste.

#### Dendrite and axon injury assays

Imaging, excluding immunohistochemistry, was performed in living whole-mount larvae. For all assays involving injury, dendrites or axons were severed from da neurons by focusing a two-photon 860-900-nm laser mounted on a Zeiss LSM 780 or Zeiss LSM 980 fluorescence microscope on a dendrite using methods described previously.^14–16^ To immobilize the larva for injury, larva were mounted on a glass slide between a 4% agarose pad and a coverslip using glycerol as a mounting media. For larger larva, vacuum grease and tape was used to keep the coverslip in place and immobilize the larva. To injure dendrites or axons on the Zeiss LSM 780, neurons were imaged using the 2-photon and cut with the 2-photon by focusing the laser on the desired area until bleaching of the fluorophore signal was seen, indicating a cut. To injure on the Zeiss LSM 980, neurons were imaged using 488 nm or 560 nm green or red lasers and injured using the bleaching function with the 2-photon. Axons were distally severed ∼40 uM away from the cell body to avoid dendrite-axon conversion.^106^ For the single branch injury assay, the laser was focused on a single primary dendrite branch near the cell body. For the terminal branch injury assay, the laser was focused on a terminal dendrite branch near the cell body. For the near branch injury assay, the laser was focused to a point near a dendrite branch for a comparable amount of time to dendrite injury assays. Neurons were imaged on a Zeiss LSM 700 or Zeiss LSM 980 fluorescence microscope 24 hours after injury to confirm successful injury and, for some experiments, at 72 hours after injury to assess regeneration. Uninjured control neurons were imaged on the day of injury but left alone and then imaged at 24 and 72 hours after injury for comparison to injured neurons. All injury assays were performed such that there was sufficient time for regeneration to occur after injury before pupation (at least 72 hours).

#### Immunohistochemistry

Larva were anesthetized with either ice or isoflurane then fileted and fixed for 30 minutes in 4% paraformaldehyde at room temperature after the final imaging time point for these experiments. Following fixation, larval filets were rinsed and washed with PBS with 1% Triton-X then incubated for 1 hour and 30 minutes in blocking buffer (PBS/0.5% TritonX/10% Horse Serum). Following blocking, filets were incubated with primary antibodies (Mouse anti-futsch 1:2, rabbit anti-GFP 1:1000) overnight at 4°C. Following primary antibody incubation, filets were rinsed and then washed in PBS with 0.5% Triton-X 5 times for 10 minutes each. Filets were then incubated with secondary antibodies (Goat anti-mouse 647 and Goat anti-rabbit 488) overnight at 4°C. Filets were then rinsed and washed in PBS with 0.5% Triton-X 5 times for 15 minutes each.

Filets were mounted on glass slides with 1:1 1xPBS and Glycerol for later imaging.

### QUANTIFICATION AND STATISTICAL ANALYSIS

Dendrite arbors were traced using the Simple Neurite Tracer plugin in ImageJ to determine the number of dendrite branch tips and the total length of all the dendrite branches. For some neurons, using these traced arbors, Sholl analysis of dendrite branches crossing circles separated by 1 μm was performed. For single branch injury analysis, normalized control neurons were used for comparison. Normalized controls represent uninjured neurons normalized for one less primary dendrite branch by eliminating the least branched primary dendrite branch and its secondary and terminal branches from quantification. For all graphs comparing means, values are plotted as mean ± SEM and individual data points representing individual neurons are shown. For paired data, values are plotted for individual neurons with faded lines connecting repeated measurements at 24 and 72 hours AI. Solid lines represent the slope (m) between the mean 24 and 72 hour AI values which were calculated with the formula. Percent branches gained and percent length gained were calculated with the formula *m* =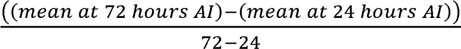. For both GMA and TauGFP analysis, mean RFU values for branches were calculated in ImageJ by measuring the mean fluorescence of ∼50 μm stretches of individual primary branches (starting as close to the cell body as possible) and subtracting local mean background fluorescence around the branch and then calculating the mean RFU of all primary branches taken together. For both GMA and TauGFP analysis, median RFU for neuron cell bodies were calculated in ImageJ by measuring the median fluorescence of the entire cell body and subtracting local median background fluorescence around the cell body. For EB-1 comet analysis, we used the open-source version of the software KymoButler to measure comet velocity.^107^

Sample sizes are represented within figures other than the following. Figure 1B, 1C, Supp1D, Supp 1C: WT 24-168 hrs AEL n= 14, n=10, n=10, n=10, n=9, n=5, n=5 respectively; ATX1.82Q 24-168 hrs AEL n=9, n=10, n=10, n=10, n=8, n=9, n=6 respectively; ATX2.64Q 24-168 hrs AEL n=10, n=6, n=8, n=11, n=11, n=7, n= 10 respectively; MJD.78Q 24-168 hrs AEL n=10, n=9, n=12, n=7, n=8, n=11, n=10 respectively, and HTT231NT.128Q 24-168 hrs AEL n=12, n=6, n=5, n=5, n=5, n=9, n=7 respectively. Figure Supp1B: WT 48 hrs AEL n=1, WT 96 hrs AEL n=3 WT 120 hrs AEL n=2 ATX1.82Q 48 hrs AEL n=1, ATX1.82Q 96 hrs AEL n=3, ATX1.82Q 120 hrs AEL n=2, MJD.78Q 48 hrs AEL n=4, MJD.78Q 96 hrs AEL n=3, MJD.78Q 120 hrs AEL n=2. Figure Supp1G: WT 24-120 hrs AEL n=3, n=3, n=3, n=5, n=15 respectively; MJD.78Q 24-120 hrs AEL n=6, n=4, n=3, n=5, n=7 respectively; ATX1.82Q 120 hrs AEL n=15. Figure Supp5E: WT UI n=5, WT SBI n=5, ATX1.82Q UI n=7, ATX1.82Q SBI n=7, ATX2.64Q UI n=6, ATX2.64Q SBI n=4.

Statistical analysis was performed using Graphpad Prism software and Microsoft Excel software. The statistical details of experiments can be found in the figure legends. Statistical significance was tested using the two-tailed Welch’s t-test for all analysis comparing two groups, using paired t-test for all analysis comparing the same neuron at two time points, and using one-way ANOVA with either Dunnett’s or Tukey’s multiple comparisons correction for analysis comparing more than two groups. For all statistical tests used ****p < 0.0001, ***p < 0.001, **p < 0.01, *p < 0.05, ^ns^p ≥ 0.05.

## SUPPLEMENTAL INFORMATION TITLES AND LEGENDS

**Supplemental Figure 1.**
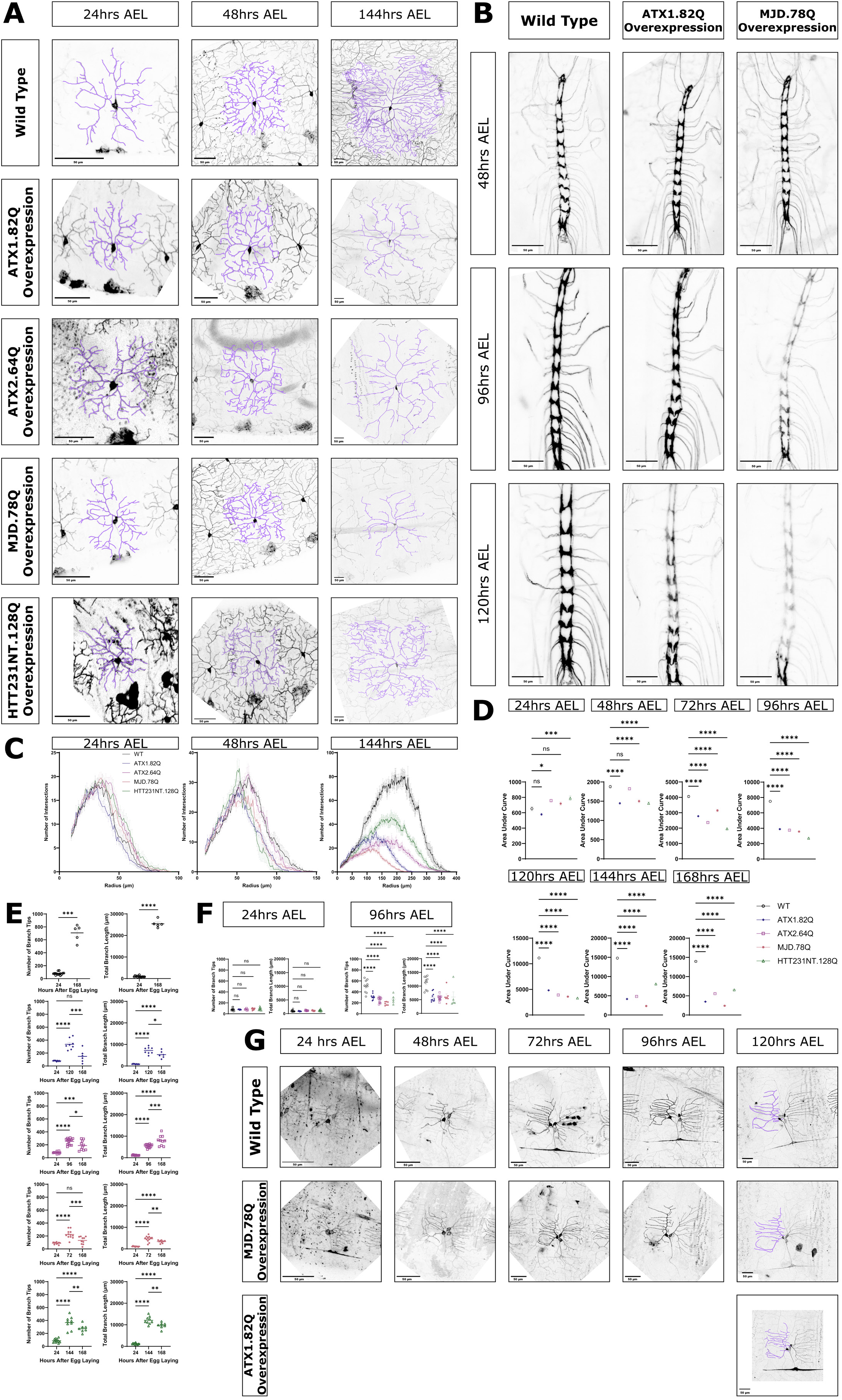
Class IV neurons overexpressing pathogenic polyglutamine proteins experience progressive degeneration of dendrites. A) WT neurons and neurons overexpressing ATX1.82Q, ATX2.64Q, MJD.78Q, and HTT231NT.128Q at 24, 48, and 144 hrs AEL. Scale bar 50 μm. B) Ventral nerve cord for WT neurons, and neurons overexpressing ATX1.82Q and MJD.78Q at 48, 96, and 120 hrs AEL. Scale bar 50 μm. C) Sholl analysis at 24, 48, and 144 hrs AEL for WT neurons, and neurons overexpressing ATX1.82Q, ATX2.64Q, MJD.78Q, and HTT231NT.128Q. Legend applies for all graphs in C. D) Area under the Sholl curve at 24-168 hrs AEL for WT neurons and neurons overexpressing ATX1.82Q, ATX2.64Q, MJD.78Q, and HTT231NT.128Q. Legend applies for all graphs in D, E, and F. Mean ± SEM. One-way ANOVA with Dunnett’s multiple comparisons correction. E) Starting and endpoint values for dendrite branches and length for WT neurons (top). Welch’s t-test. Starting, maximum, and endpoint values for dendrite branches and length for ATX1.82Q neurons (second from top), ATX2.64Q neurons (third from top), MJD.78Q neurons (fourth from top), and HTT231NT.128Q neurons (bottom). One-way ANOVA with Tukey’s multiple comparisons correction. F) Dendrite branches and length at 24 hrs AEL and 96 hrs AEL between WT neurons and ATX1.82Q, ATX2.64Q, MJD.78Q, and HTT231NT.128Q neurons. One-way ANOVA with Dunnett’s multiple comparisons correction. G) Class 1 ddaE WT and MJD.78Q overexpression neurons at 24-120 hrs AEL and ATX1.82Q overexpression neurons at 120 hrs AEL. Scale bar 50 μm.

**Supplemental Figure 2.**
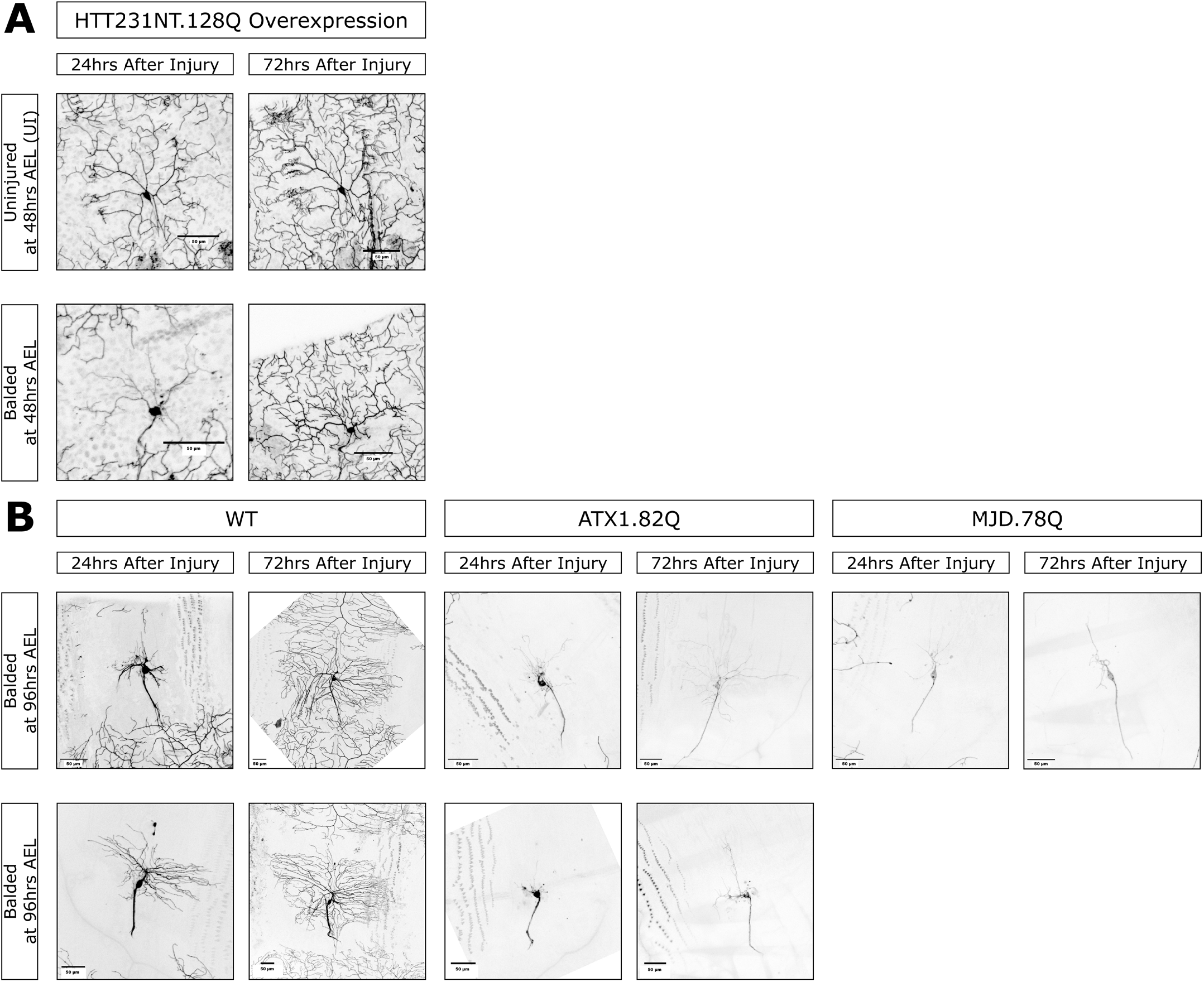
Pathogenic polyglutamine protein expression does not prevent dendrite regeneration. A) Uninjured (top) and balded at 48 hrs AEL (bottom) HTT231NT.128Q overexpression neurons at 24 and 72 hrs AI. Scale bar 50 μm. B) WT (left), ATX1.82Q overexpression (middle), and MJD.78Q overexpression (right) neurons balded at 96 hrs AEL. Scale bar 50 μm.

**Supplemental Figure 3.**
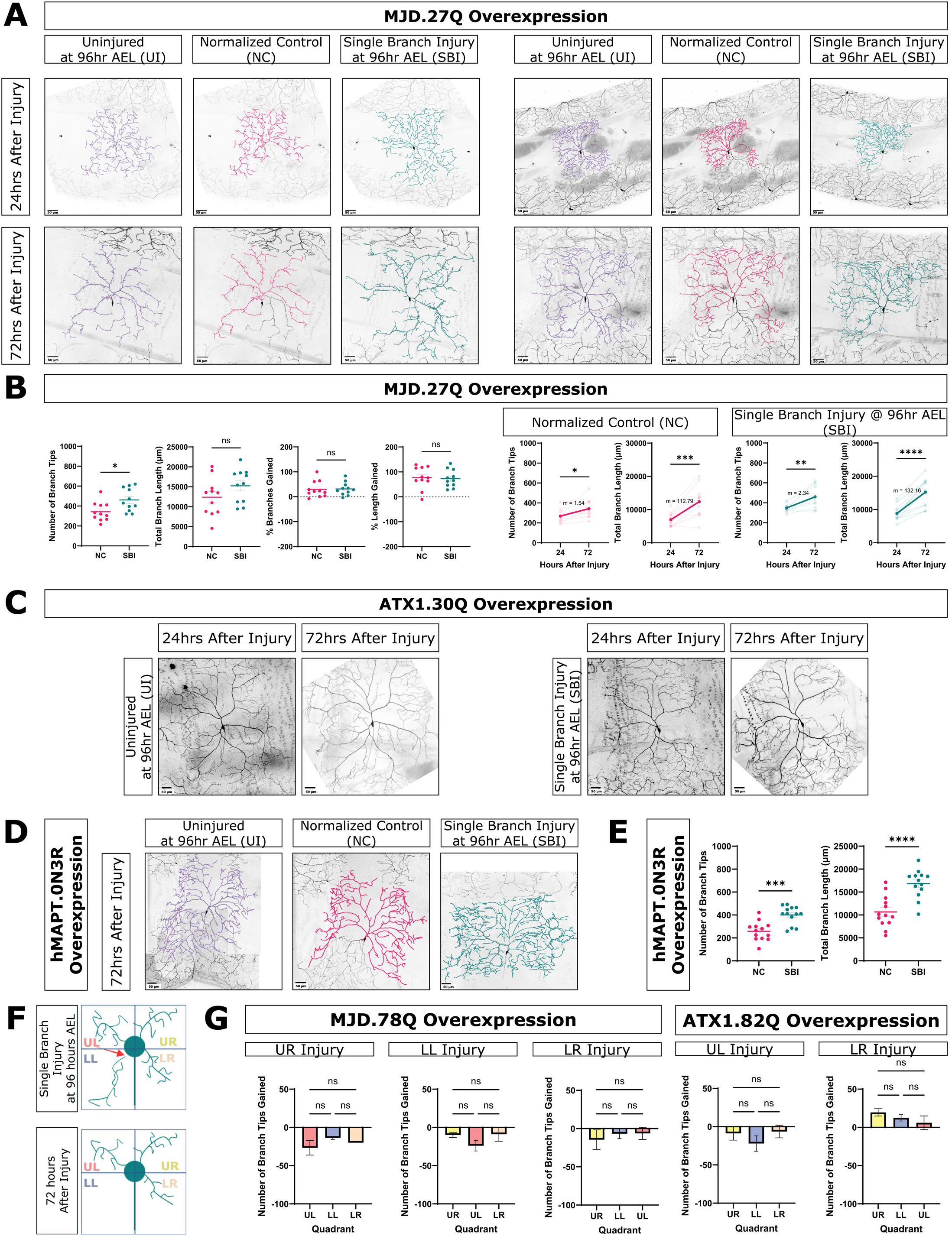
Short poly-Q neurons respond to SBI like WT neurons, SBI induces neuroprotection in hMAPT overexpression neurons, and location of SBI does not guide new growth. A) Uninjured neurons, normalized control neurons, and single branch injured neurons at 24 and 72 hrs after injury for two different animals with class IV neurons overexpressing MJD.27Q. Scale bar 50 μm. B) Number of branch tips and total branch length of NC and SBI neurons at 72 hrs AI or comparing % change in number of branch tips and total branch length between 24 to 72 hrs AI between NC and SBI neurons. Welch’s t-test. Number of branch tips and total branch length at 24 and 72 hrs AI for NC (left, pink lines) and SBI (right, green lines) MJD.27Q neurons. Paired t-test. C) Uninjured and SBI ATX1.30Q overexpression neurons at 24 and 72 hrs after injury. Scale bar 50 μm. D) Uninjured, normalized control, and single branch injured neurons at 72hr AI for hMAPT.0N3R overexpression neurons. E) Number of branch tips and total branch length for NC and SBI hMAPT.03NR overexpression neurons at 72 hrs AI. F) Schematic demonstrating quadrants for analysis of arbor growth location following SBI. UL = upper left, LL = lower left, UR = upper right, LR = lower right. E) Number of branch tips gained for different quadrants of the dendrite arbor following injury to a branch in another quadrant for MJD.78Q and ATX1.82Q overexpression neurons. One-way ANOVA with Tukey’s multiple comparisons correction.

**Supplemental Figure 5.**
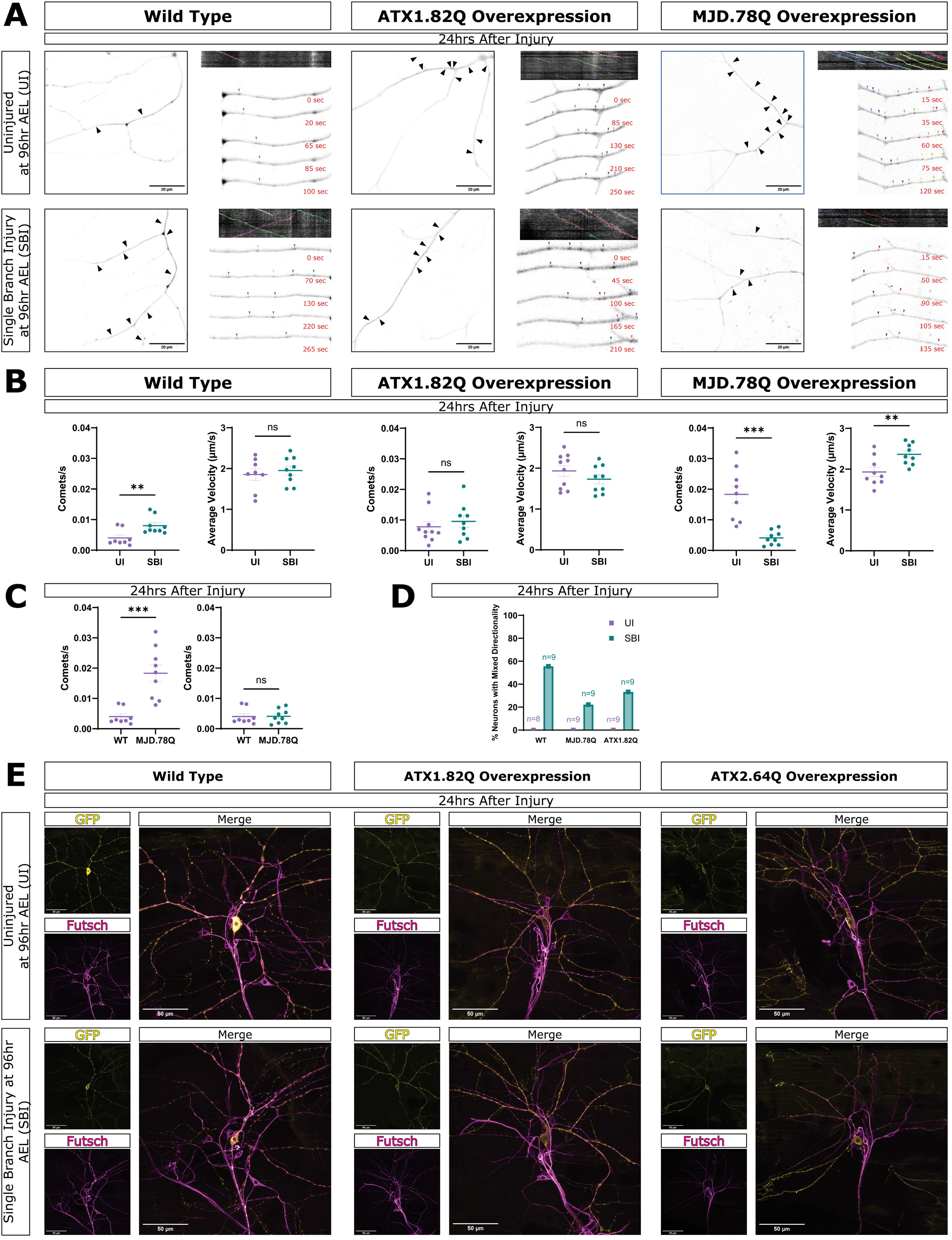
Neither microtubule composition nor dynamics are rescued in single branch injured polyQ neurons. A) Snapshots of EB-1 comet tracking movies for uninjured and single branch injured WT and ATX1.82Q and MJD.78Q overexpression neurons at 24 hrs after injury. Arrows indicate individual EB-1 comets and colored arrows mark the same comet at different time points. Scale bar 20 μm. B) Comets/s for a 10 μm length of dendrite and average comet velocity between UI and SBI neurons for WT and ATX1.82Q and MJD.78Q overexpression neurons at 24 hrs after injury. Welch’s t-test. C) Comparison of comets/s for a 10 μm length of dendrite between uninjured WT (purple) and uninjured MJD.78Q neurons (purple) and comparison of comets/s for a 10 μm length of dendrite between uninjured WT (purple) and SBI MJD.78Q overexpression neurons (green) at 24 hrs after injury. Welch’s t-test. D) % neurons with mixed EB-1 comet directionality (retrograde, anterograde) at 24 hrs after injury for WT and ATX1.82Q and MJD.78Q overexpression neurons. Values are plotted as percent of all movies per injury type and genotype. Sample size n represents the total number of neuron movies for each injury type and genotype. E) Immunostaining for GFP (pseudo-color yellow) and futsch (pseudo-color magenta) of uninjured and single branch injured WT and ATX1.82Q and ATX2.64Q overexpression neurons.

